# Motor unit dysregulation following 15 days of unilateral lower limb immobilisation

**DOI:** 10.1101/2022.06.01.494421

**Authors:** Thomas B. Inns, Joseph J. Bass, Edward J.O. Hardy, Daniel J. Wilkinson, Daniel W. Stashuk, Philip J. Atherton, Bethan E. Phillips, Mathew Piasecki

## Abstract

Disuse atrophy, caused by situations of unloading such as limb immobilisation, causes a rapid yet diverging reduction in skeletal muscle function compared to muscle mass. While mechanistic insight into the loss of mass is well studied, deterioration of muscle function with a focus towards the neural input to muscle remains underexplored. This study aimed to determine the role of motor unit adaptation in disuse-induced neuromuscular deficits. Ten young, healthy male volunteers underwent 15 days of unilateral lower limb immobilisation with intramuscular EMG (iEMG) recorded from the vastus lateralis during knee extensor contractions normalised to maximal voluntary contraction (MVC) pre and post disuse. Muscle cross-sectional area was determined by ultrasound. Individual MUs were sampled and analysed for changes in MU discharge and MU potential (MUP) characteristics. Vastus lateralis (VL) CSA was reduced by approximately 15% which was exceeded by a two-fold decrease of 31% in muscle strength in the immobilised limb, with no change in either parameter in the non-immobilised VL. Parameters of MUP size were largely reduced with immobilisation, while neuromuscular junction (NMJ) transmission instability increased, and MU firing rate decreased at several contraction levels. All adaptations were observed in the immobilised limb only. These findings highlight impaired neural input following immobilisation reflected by suppressed MU discharge rate and instability of transmission at the NMJ which may underpin the disproportionate reductions of strength relative to muscle size.

**Key points:**

- Muscle mass and function decline rapidly in situations of disuse such as bed rest and limb immobilisation.
- The reduction in muscle function commonly exceeds that of muscle mass, which may be associated with the dysregulation of neural input to the muscle.
- We have used intramuscular electromyography to sample individual motor unit and near fibre potentials from the vastus lateralis following 15 days of unilateral limb immobilisation. Following disuse, the disproportionate loss of muscle strength when compared to size was associated with suppressed motor unit firing rate and increased markers of neuromuscular junction transmission instability.
- These central and peripheral motor unit adaptations were observed at multiple contraction levels and in the immobilised limb only. Our findings demonstrate neural dysregulation as a key component of functional loss following muscle disuse in humans.

## Introduction

Disuse atrophy is the loss of skeletal muscle mass associated with decreased external loading or complete immobilisation. It is common in clinical settings following joint trauma, nerve injury, or prescribed bed rest, and progresses rapidly with reductions of strength occurring after just 5 days (Wall *et al*., 2014). As such, it has been widely applied as an experimental model to investigate underpinning mechanisms in scenarios of space flight, prolonged bed rest, spinal cord injury, and ageing (Castro *et al*., 1999; Narici & De Boer, 2011; Puthucheary *et al*., 2013). The trajectory of decline is not linear; five days of unilateral lower limb suspension lead to ∼3 % reduction in quadricep cross-sectional area (CSA) (Wall *et al*., 2014) and eight weeks of bed rest reduced quadriceps CSA by ∼14% (Mulder *et al*., 2006).

The loss of muscle strength with disuse is commonly reported to exceed the loss of muscle size. Following 8 weeks of bedrest, quadriceps CSA declined by 14% compared to a 17% decline in strength (Mulder *et al*., 2006), while only 10 days of bed rest was enough to elicit a ∼6% reduction in CSA and a ∼14% reduction in knee extensor maximal voluntary contraction (MVC) (Monti *et al*., 2021). Furthermore, meta-analysis of bed rest studies with a combined 118 participants across various durations of disuse found no relationship between length of bed rest and muscle size, whereas muscle power reduction and bed rest duration were strongly related (Di Girolamo *et al*., 2021). Numerous data provide mechanistic insight to muscle *atrophy*, including decreased MPS, increased MPB, mitochondrial dysfunction, insulin resistance, and histochemical markers of fibre denervation (Phillips *et al*., 2009; Rudrappa *et al*., 2016; Monti *et al*., 2021), yet mechanistic insight into loss of muscle *function* is less clear.

Successful activation of muscle relies upon coordinated input to the motoneuron pool and synaptic transmission across neuromuscular junctions (NMJs), with increases in muscle force mediated by the recruitment and rate modulation of functioning motor units (MUs). Motor units from hand muscles showed a marked reduction in FR following disuse, which corresponded with reduced force generating capacity (Duchateau & Hainaut, 1990). Biopsies from the VL following 10 days of bed rest revealed structural disruption at the NMJ (Monti *et al*., 2021) suggesting a contribution to decreased muscle activation and function. Furthermore, electromyography (EMG) amplitude following 14 days of unilateral lower limb suspension (ULLS) decreased in the immobilised limb along with peak torque, while limb mass remained unchanged (Deschenes *et al*., 2002), all of which highlight potential neural dysregulation. The extent of disuse atrophy is also muscle dependent which has led to the descriptors of ‘atrophy resistant’ and ‘atrophy susceptible’ muscles (Bass *et al*., 2021). A recent systematic review consolidated previous work to date (Campbell *et al*., 2019), and of the 40 studies included, 20 immobilised the knee, preventing use of one of the most functionally relevant muscle groups, i.e., the quadriceps. Furthermore, just 20 studies investigated neural factors (i.e., using surface EMG), yet adaptations of individual MU characteristics, particularly at the NMJ, remain largely unexplored.

The purpose of the study was to quantify neuromuscular changes by studying individual MU features in the immobilised and non-immobilised limbs by utilising intramuscular EMG (iEMG) to sample individual MUs from the VL pre and post 15-days of immobilisation. We hypothesised that, coinciding with a reduction in muscle mass and strength, NMJ transmission instability would increase, and MU firing rate would decrease in the immobilised leg, with no change to the non-immobilised limb.

## Methods

### Ethical approval

This study was approved by the University of Nottingham Faculty of Medicine and Health Sciences Research Ethics Committee (103-1809) and conformed with the Declaration of Helsinki. It was registered online at clinicaltrials.gov (NCT04199923). Participants were recruited locally from the community via advertisement posters in print and on research group social media pages. Ten healthy, young male participants were recruited to take part in the study. After providing written informed consent to participate in the study, potential participants were screened for eligibility against pre-determined exclusion criteria, including active cardiovascular, cerebrovascular, respiratory, renal, or metabolic disease, active malignancy, musculoskeletal or neurological disorders. Once eligibility was confirmed, participants were invited to the laboratory for baseline testing, as described below.

### Muscle ultrasound

Vastus lateralis (VL) cross sectional area scans (CSA) were taken at the mid-belly of the muscle. This was located by measuring the length from the midline of the patella to the greater trochanter and taking the middle value of that line. Following this, a narrow ultrasound probe (LA523 probe and MyLab™50 scanner, Esaote, Genoa, Italy) was used to find the medial and proximal borders of the muscle where the aponeurosis of the VL intersected with the *vastus intermedius*. Three axial plane images were collected following this line from both legs and subsequently analysed using ImageJ (Laboratory of Optical and Communication, University of Wisconsin-Madison, WI, USA) to quantify CSA (Scott *et al*., 2017).

### Lower limb power

Explosive unilateral lower limb power was assessed using the Nottingham Power Rig (University of Nottingham; (Bassey & Short, 1990)). For this, participants were seated on the purpose-built rig with their knee at 90° when the foot was on the plate. The rig is designed to isolate power production from the lower limb. Following a 3 s countdown, participants were instructed to perform an explosive push. The power exerted was displayed digitally and the highest of three attempts recorded.

### Maximal voluntary isometric contraction

Participants were seated in a custom-built isometric dynamometer with their knee joint angle fixed at 90° while the hip joint angle was at 110°. The ankle was secured in place to a plate connected to a force transducer. Three moderate intensity warm-up contractions were carried out with visual feedback of force traces on a screen in front of the participants. Using a waist belt to prevent hip lifting and facilitate isolation of the knee extensors, participants were verbally encouraged to perform an isometric knee extension at maximal capacity.

### Vastus lateralis motor point identification

Using low intensity percutaneous electrical stimulation (400V, pulse width 50 µS, current ∼10 mA; delivered via a Digitimer DS7A, Welwyn Garden City, UK), the surface of the VL was explored to find the point producing the greatest visible twitch with the lowest current, i.e., the motor point (Piasecki *et al*., 2016*a*).

### Force and electromyography recording

Surface electromyography was arranged in a bipolar configuration. The recording electrode was placed over the identified motor point and the reference electrode over the patellar tendon (disposable self-adhering Ag-AgCl electrodes; 95 mm^2^; Ambu Neuroline, Baltorpbakken, Ballerup, Denmark). A ground electrode was placed just above the reference electrode on the patella (Ambu Neuroline Ground). EMG signals were digitised (CED Micro 1401; Cambridge Electronic Design, Cambridge, UK) and Spike2 (version 9.00a, CED) software was used to provide a real-time display of the signal on screen. Peak twitch force was calculated using percutaneous electrical stimulation (Digitimer, UK), and the maximal M-wave was measured with surface electromyography at the motor point. A stimulating pen was placed over the femoral nerve in the inguinal fold and stimulation intensity (400V, pulse width 50 µS, current typically 70-110 mA) was increased until the M-wave amplitude displayed in Spike2 plateaued. The peak force generated corresponding to the maximal M-wave was recorded, and all force signals were recorded at 100 Hz. Force steadiness was assessed during the sustained voluntary contractions as the coefficient of variation of the force deviation from the target line, averaged at each contraction intensity. The first two passes of the target (<1s) were excluded from the calculation to avoid corrective actions when reaching the target line.

### Intramuscular electromyography

A concentric needle electrode (Ambu Neuroline model 740 25-45/25, Ambu, UK) was inserted at the *vastus lateralis* motor point. The recorded iEMG signals were sampled at 50 kHz and bandpass filtered from 10 Hz to 10 kHz (1902 amplifier, CED). A real-time display was observed using Spike2 (CED) and data were stored offline for analysis. Once the needle was inserted, contractions were carried out at 10% and 25% of the participants MVC following a visual target line. Six contractions were recorded at each intensity, with the needle electrode position altered between each to sample a broader range of MUPs (Jones *et al*., 2021). From the original position, the needle electrode was slightly withdrawn, and the bevel rotated 180° at each depth and positioned to maximise signal to noise ratio. These voluntary contractions were held for 12 s with a 10-20 s rest in between each contraction.

### EMG analysis

Decomposition-based quantitative electromyography (DQEMG) software was used to process raw EMG data. This involved the detection of MUPs and extraction of MUP trains (MUPTs) from individual MUs generated during sustained iEMG signals recorded during voluntary contractions at set intensities. MUPTs were excluded if they contained MUPs from multiple MUs or fewer than 40 MUPs. MUP templates were visually inspected to ensure markers were placed correctly to start, end, positive and negative peaks of the waveforms. MUP area was defined as the integral of the absolute value of MUP values recorded between start and end markers multiplied by the sampling time interval in μV/ms (Piasecki *et al*., 2021*b*). MUP amplitude was determined as the measurement from the maximal positive and negative peaks of the waveform (Guo *et al*., 2022). MUP complexity, measured as the number of turns, was defined as the number of significant slope direction changes within the duration of the MUP of a height >20 μV. A near fibre MUP (NF-MUP) was obtained from each MUP by estimating the slopes of each MUP (Piasecki *et al*., 2021*b*) and NMJ transmission instability, measured as near-NF-MUP jiggle, was determined as the normalised means of median consecutive amplitude differences (Piasecki *et al*., 2021*b*). MU firing rate (MU FR) was recorded as the rate of occurrence per second of MUPs within a MUPT in Hz, and MU FR variability was determined as the coefficient of variation of the inter-discharge interval.

### Immobilisation procedure

For the 15 days of immobilisation, a unilateral lower-limb suspension (ULLS) model was used. The knee joint was fixed at 75° flexion using a hinged leg brace (Knee Post op Cool, Össur, Iceland) with the ankle joint fixed using an air-boot (Rebound Air Walker, Össur), ensuring that the immobilised leg was not able to bear any weight. Crutches were provided and adjusted according to the height of the participant, and training on their effective use was provided. The brace and boot remained in place at all times, including sleeping and bathing, with tamper tags attached to each to monitor intervention adherence.

### Follow-up testing

Following 15 days of ULLS, participants were invited back to the laboratory for post-immobilisation testing. All procedures carried out in the baseline testing visit were repeated in both legs. In addition to measuring FS and iEMG at 25% of post-immobilisation MVC (i.e., relative force, referred to as follow-up force), iEMG was also performed at 25% of pre-immobilisation MVC (i.e., absolute force, referred to as baseline force) to compare FS and MU characteristics normalised to pre and post disuse-induced strength loss.

### Statistical analysis

Statistical analysis of MVC and ultrasound CSA was carried out using GraphPad Prism version 9.1.0 (GraphPad Software, CA, USA) using repeated measures 2-way analysis of variance with Šidak’s post-hoc analysis. Multi-level mixed effect linear regression models were used to analyse MU parameters, in StataSE (v16.0, StataCorp LLC, TX, USA). For both analyses, two within-subject factors were assessed: time (pre and post) and leg (immobilised and control). Where no leg x time interactions were present, separate models were performed to assess effects of time in individual limbs. Additional exploratory analyses were performed to investigate relationships between iEMG variables and key physiological outcome variables – muscle strength (MVC) and size (CSA) using R (Version 4.2.0, (https://cran.r-project.org/) implemented using R studio. Firstly, correlative analysis was assessed with Pearson’s product moment correlation coefficient and visualised using corrplot (https://cran.r-project.org/web/packages/corrplot) to determine any strong relationships between variables. Cluster analysis and principal component analysis for variables were performed using the ClustOfVar (https://cran.r-project.org/web/packages/ClustOfVar) and factoextra (https://cran.r-project.org/web/packages/factoextra) packages respectively, to determine which variables strongly clustered and related to others. Finally, using a subset of these best clustering variables, multivariate linear regression was performed to determine which clustered variables best predict changes in MVC and CSA. Significance was assumed if p<0.05.

## Results

### Participant Characteristics

Ten male participants took part in this study. Characteristics are shown in table 1.

**Table 1:** Descriptive characteristics of participants showing mean and standard deviation (SD)

| N = 10 | Mean (SD) |
| --- | --- |
| Age (years) | 23.68 (3.36) |
| Height (cm) | 181.56 (6.69) |
| Weight (kg) | 79.37 (9.70) |
| BMI (kg/m<sup>2</sup>) | 24.02 (2.10) |

### Muscle Size and Function

There was a significant leg x time interaction in VL CSA (p<0.001, n=9), which decreased in the immobilised leg after 15 days (−15%, p<0.001) and remained unchanged in the control leg (p=0.78, fig. 2A). Similarly, there was a significant leg x time interaction of knee extensor MVC (p<0.01), which decreased in the immobilised leg (−31%, p<0.001) while no change was seen in the control leg (p=0.498, fig. 2B). Peak twitch force (n=7) showed no significant leg x time interaction (p=0.0582). No change was seen in either the immobilised leg (p=0.130) or the control leg (p=0.410) following the immobilisation period (fig. 2C). Unilateral lower limb power output (n=9) presented a significant leg x time interaction (p=0.0172), which reduced significantly in the immobilised limb (−26%, p=0.0037) while remaining unchanged in the control leg (p=0.939, fig. 2D). Force steadiness (FS) at 10% MVC presented a significant leg x time interaction (p=0.0201, fig. 2E), with non-significant changes in the immobilised limb (−14%, p=0.065) and the control (+4%, p=0.278). No interaction was present in FS at 25% MVC (p=0.255), with no changes observed when exploring limbs individually (immobilised: p=0.999, control: p=0.221, fig. 2E).

**Figure 1:**
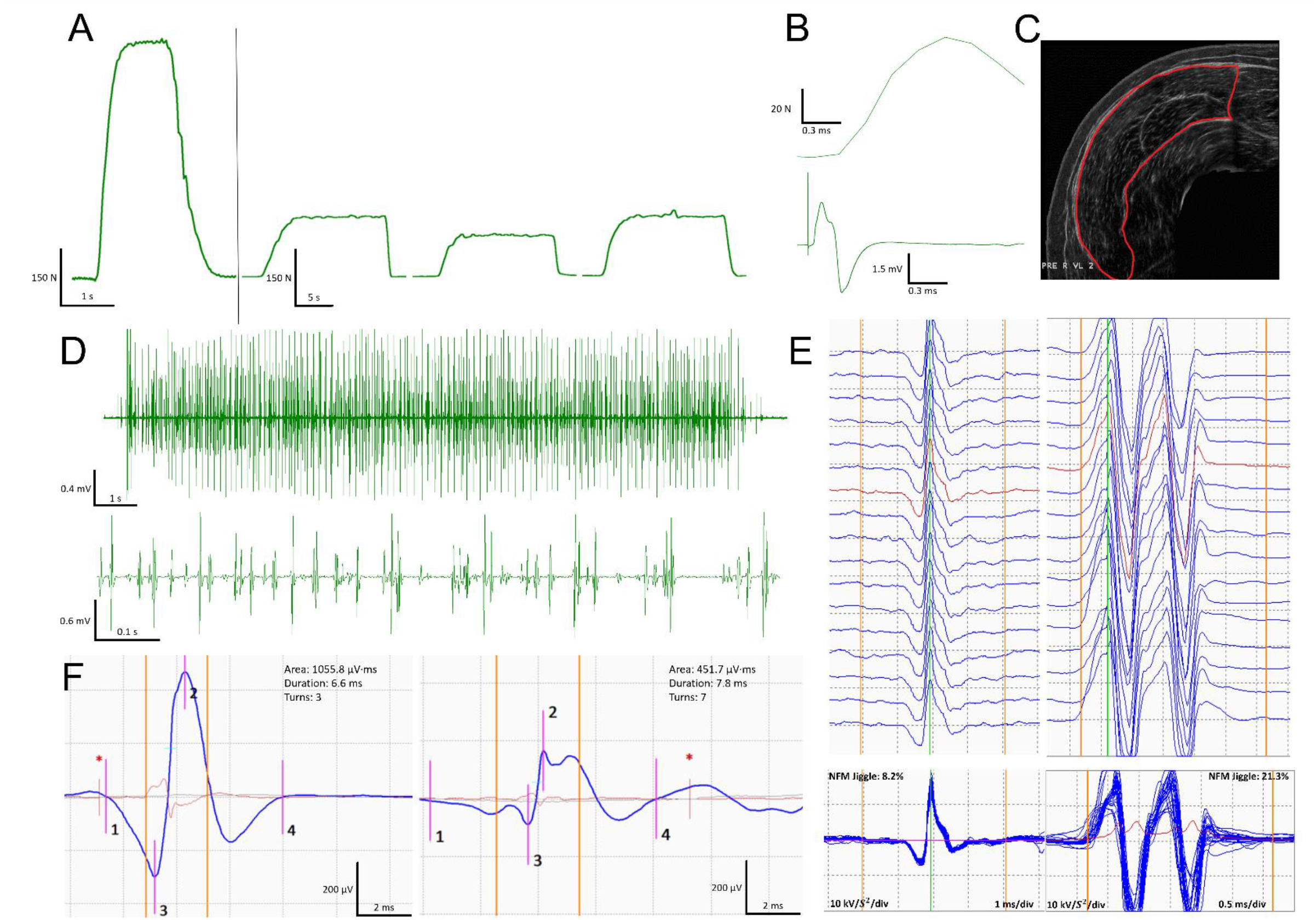
**(A)** Knee extensor force traces (left to right) from a maximal voluntary isometric contraction (MVC), 25% of MVC contraction pre 15-day unilateral limb immobilisation, 25% contraction post-immobilisation relative to follow-up MVC, and 25% contraction post-immobilisation relative to baseline MVC. **(B)** Peak twitch force recording showing force trace (upper) and corresponding surface electromyography EMG recorded M-wave (lower) from a maximal stimulation of the femoral nerve. **(C)** An example ultrasound scan of the m. vastus lateralis used to calculate cross-sectional area. **(D)** Example intramuscular EMG (iEMG) trace recorded during a 12 second voluntary isometric contraction (upper) and magnified (lower) with visible motor unit potentials (MUP). **(E)** Example raster plots (upper) and corresponding shimmer plots (lower) from near-fibre MUPs (NF-MUP) before (left) and after (right) immobilisation recorded during active contractions. **(F)** Example MUPs recorded before (left) and after (right) immobilisation. N; newtons, s; seconds, ms; milliseconds, V; volts, μV; microvolts, kV/s^2^; kilovolts per second squared, μV·ms; microvolts per millisecond.

**Figure 2:**
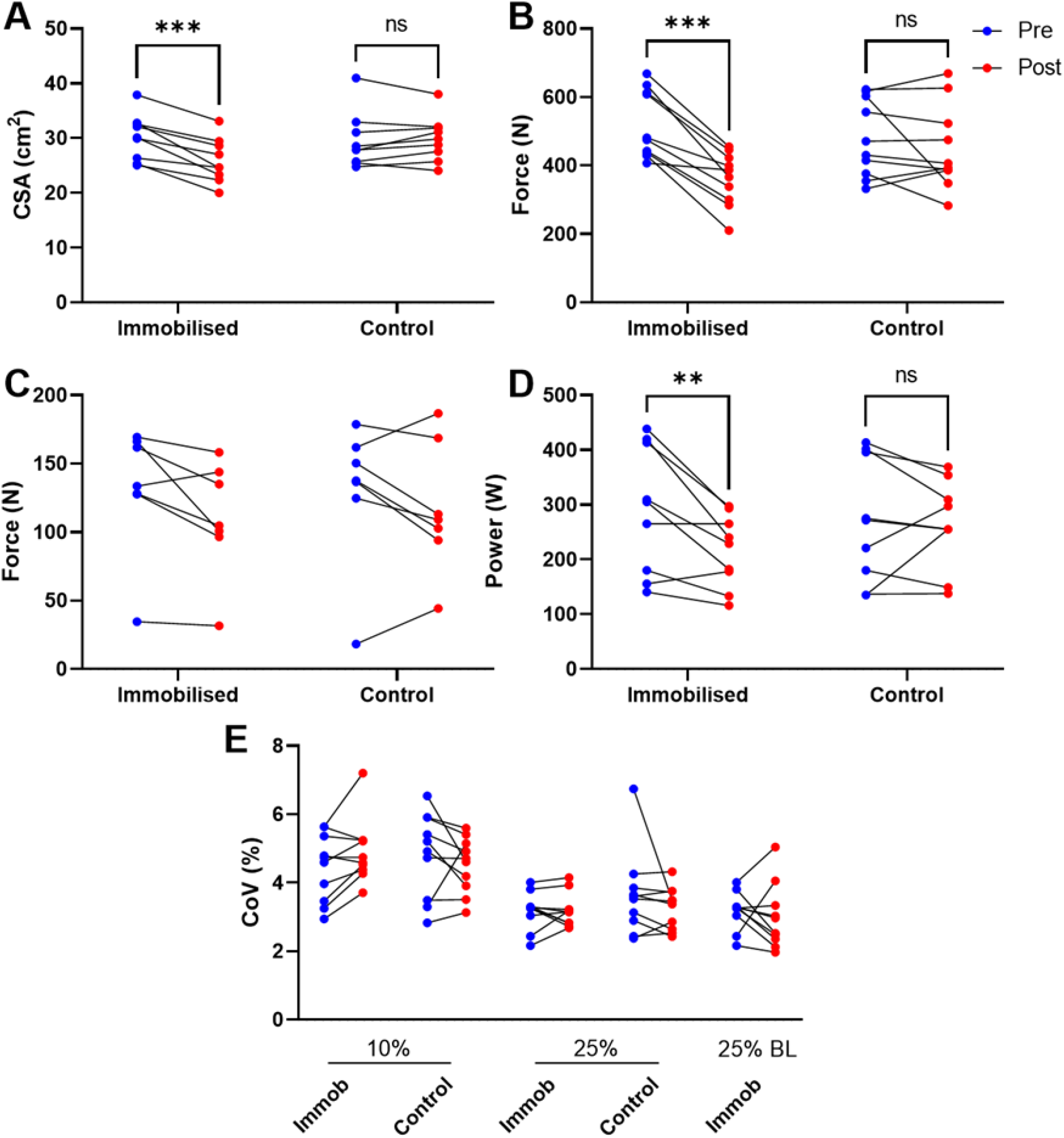
All figures show data before and after 15-day unilateral lower limb suspension in the immobilised and control legs. **A:** Vastus lateralis cross-sectional area (cm^2^) measured by ultrasound (n=9). **B:** Knee extensor maximal voluntary isometric contraction force (N). **C:** Peak twitch force (N, n=7) measured from maximal involuntary contraction elicited by femoral nerve stimulation. **D:** Unilateral lower limb power measured using the Nottingham Power Rig. **E**: Knee extensor force steadiness measured during contractions at 10 and 25% relative to both follow-up and baseline maximal voluntary isometric contraction. Coefficient of variation shown as average deviation from target line (individual mean values shown pre to post in immobilised and control legs at each contraction intensity). For all parameters, analysis performed via repeated measures 2-way analysis of variance with Šidak’s post-hoc analysis. ** = p<0.01, *** = p<0.001.

### Neuromuscular Parameters

Contractions at 25% MVC were carried out relative to follow-up MVC as well as baseline MVC. Mean values for each parameter are shown in Figures 4 and 5. MUP area at 10% MVC presented no significant leg x time interaction (p=0.091, fig. 3A). Following post-hoc analysis, a reduction was seen in the immobilised leg (p<0.001) while no change was seen in the control leg (p=0.443). At 25% follow-up MVC, a significant interaction was seen (p=0.028, fig. 4A), reflected by a reduction in the immobilised leg (p<0.001) but no change in the control (p=0.717). At 25% baseline MVC, a significant reduction in MUP area (p<0.001) was present. For MUP amplitude at 10% MVC, there was no significant leg x time interaction (p=0.106, fig. 2B). In the immobilised leg a significant reduction was present (p<0.001) with no change in the control leg (p=0.283). At 25% follow-up MVC, amplitude presented a significant leg x time interaction (p=0.046, fig. 3B), with a reduction in the immobilised leg (p=0.007) but no change in the control (p=0.881). At 25% baseline MVC MUP amplitude was also significantly reduced (p=0.006).

**Figure 3:**
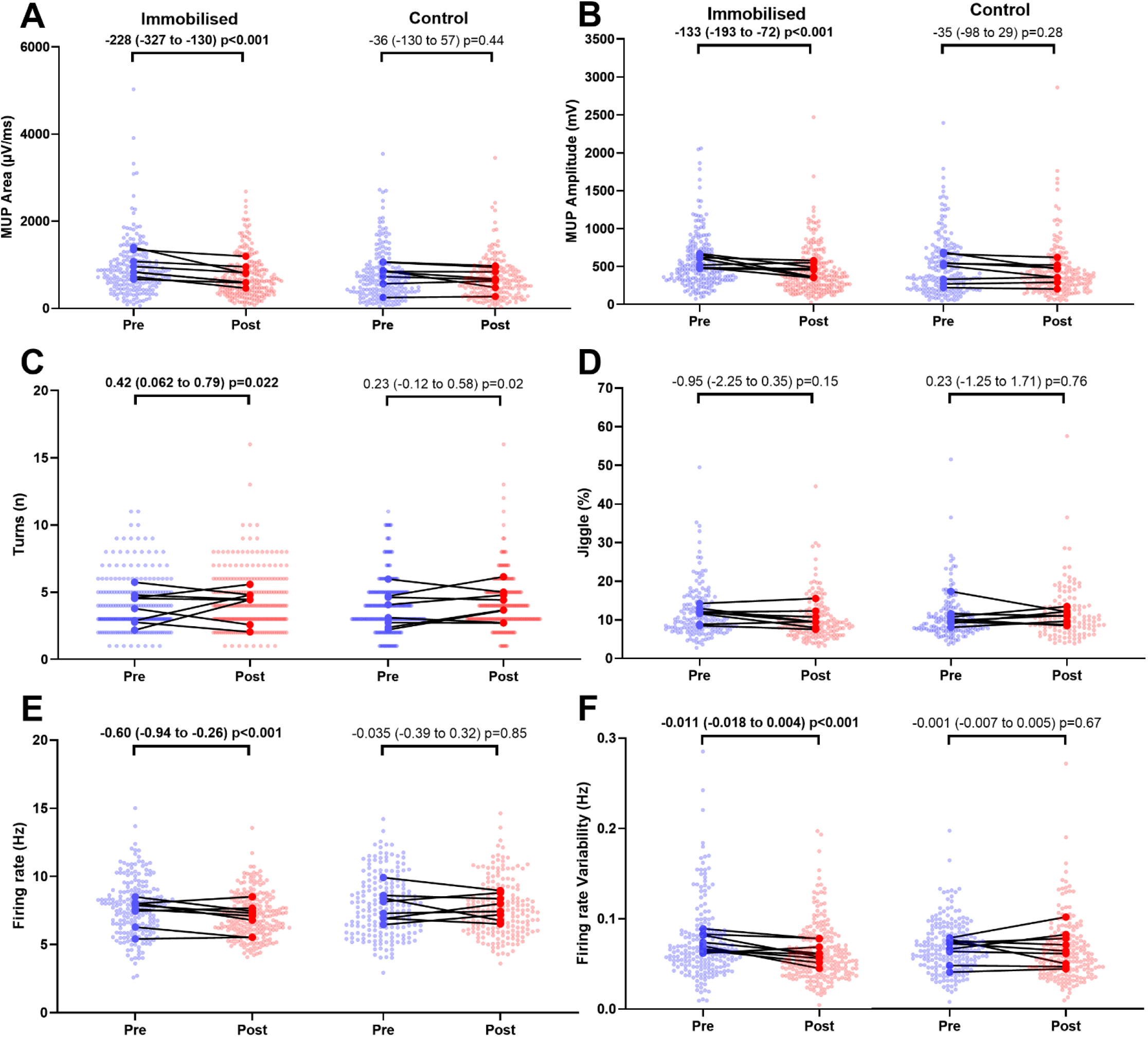
Motor unit potential parameters and MU discharge characteristics of motor units sampled during contractions performed at 10% maximal voluntary force before and after 15-day unilateral leg immobilisation in both immobilised and control legs. Data show mean values and pooled MUs with comparison bars showing β coefficient and 95% CI from multi-level mixed effects models. N = 9.

**Figure 4:**
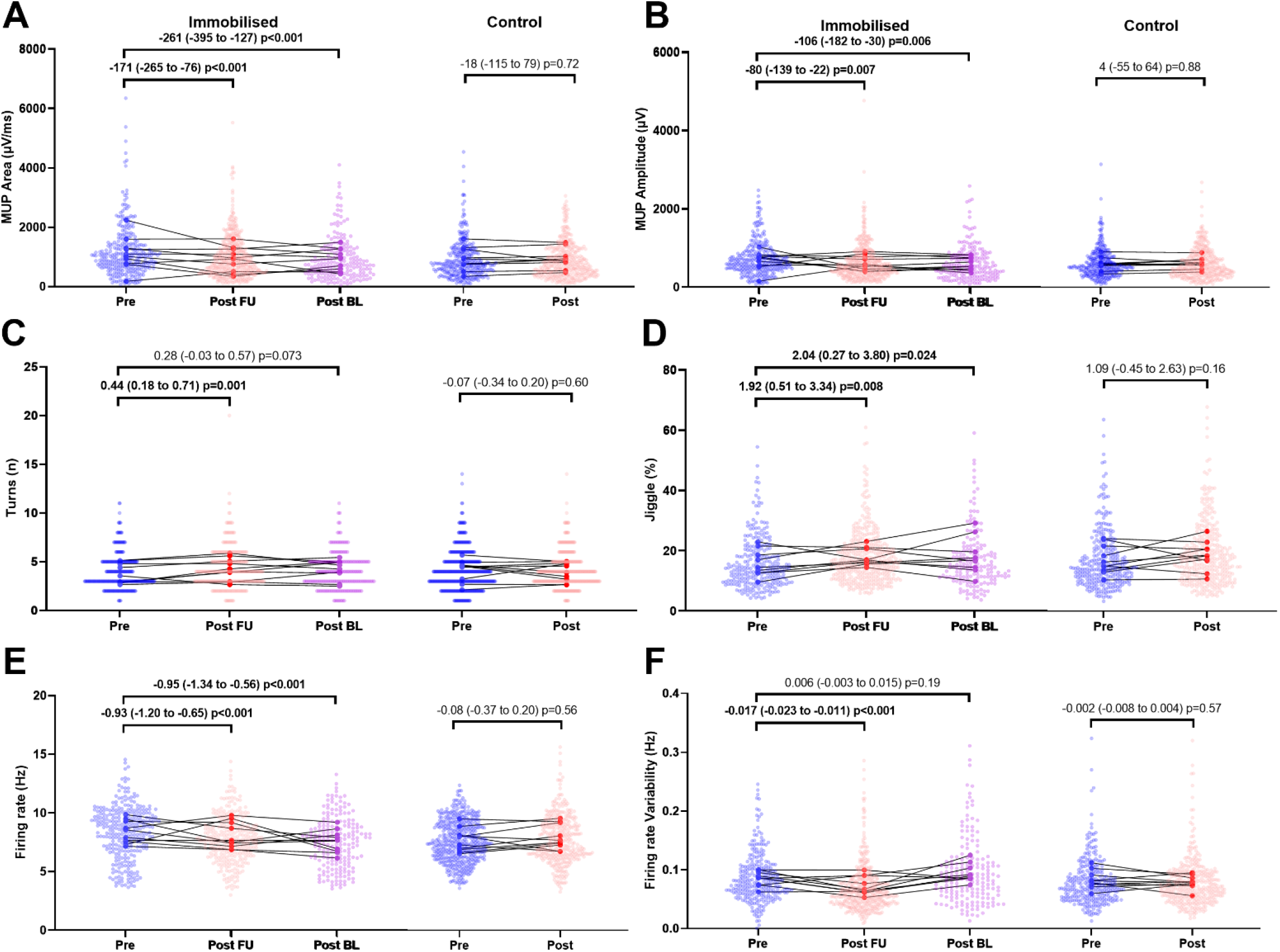
Mean values of motor unit potential parameters and MU discharge characteristics of motor units sampled during contractions performed at 25% maximal voluntary force before and after 15-day unilateral leg immobilisation in both immobilised and control legs. Values in the immobilised leg post-immobilisation are relative to follow-up MVC (column 2) and relative to baseline MVC (column 3) due to reduced MVC following immobilisation. Data show mean values and pooled MUs with comparison bars showing β coefficient and 95% CI from multi-level mixed effects models.

**Figure 5:**
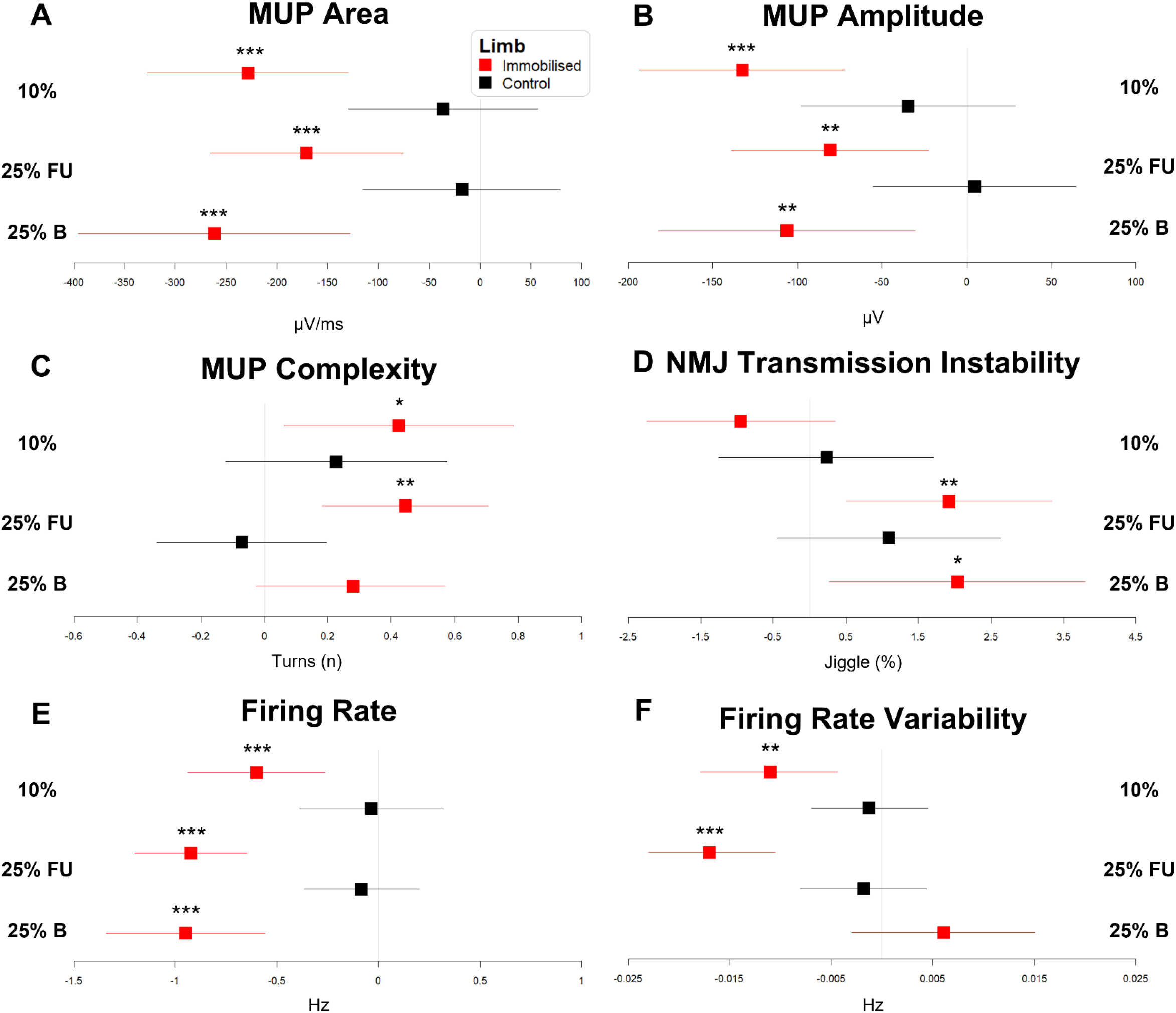
Forest plots summarising data from motor unit parameters measured from the vastus lateralis during 10% MVC, and 25% normalised to follow-up (FU) and baseline (B) MVC. Plots display beta coefficient and 95% confidence intervals from multi-level mixed linear regression models. * = p<0.05, ** = p<0.01, *** = p<0.001.

MUP complexity, defined as the number of turns, presented no significant leg x time interaction at 10% MVC (p=0.660, fig. 3C), with post-hoc analysis showing an increase in the immobilised leg (p=0.022), but no change in the control (p=0.204). At 25% follow-up MVC, complexity had a significant leg x time interaction (p=0.007, fig. 4C). An increase was seen in the immobilised leg (p=0.001) but no change was seen in the control (p=0.600). At 25% baseline MVC MUP complexity did not change significantly (p=0.073). At 10% MVC, NMJ transmission instability showed no leg x time interaction (p=0.191, fig. 3D), with no change in either the immobilised (p=0.152) or control (p=0.759) legs. Similarly at 25% follow-up MVC, no leg x time interaction was seen (p=0.966, fig. 4D). However, a significant increase was seen in the immobilised leg (p=0.008) but no change in the control (p=0.164). At 25% baseline MVC, NMJ transmission instability was also increased (p=0.024).

Motor unit FR presented a significant leg x time interaction at 10% MVC (p=0.022, fig. 3E), with a reduction in the immobilised leg (p<0.001) but no change in the control (p=0.848). At 25% follow-up MVC, an interaction was also observed (p<0.001, fig. 4E), again showing a reduction in the immobilised leg (p<0.001) but no change in the control (p=0.563). At 25% baseline MVC, firing rate was also decreased (p<0.001). Finally, firing rate variability presented no leg x time interaction at 10% (p=0.070, fig. 3F), with a reduction in the immobilised leg (p=0.001) and no change in the control (p=0.668). At 25% follow-up MVC, a significant interaction was seen (p=0.001, fig. 4F), with a reduction in the immobilised leg (p<0.001) and no change in the control (p=0.574). However, at 25% baseline MVC firing rate variability remained unchanged (p=0.193).

In exploratory analyses, to investigate potential relationships between neuromuscular parameters and muscle strength and size, correlation analysis was performed on mean differences between parameters assessed at 25% pre and 25% post MVC (fig. 6). Strong correlations were observed between MUP area and amplitude (r^2^=0.846, p=0.0003) and between NF-MUP jiggle and MVC (r^2^=0.792, p=0.0082). To more accurately reflect relationships between these variables, clustering of these variables was performed using the hierarchical clustering algorithm provided in ClustOfVar and further visualised using principal component analysis score plots (PCA, fig. 7). The key cluster of interest consisted of MVC, CSA, NF-MUP jiggle, FR and FR variability. To test whether the neuromuscular parameters in this cluster had any influence on the changes observed in muscle strength and size, the values were first normalised to the same dynamic range before multivariate linear regression was performed. There were no significant relationships for either MVC or CSA with NF-MUP jiggle (p=0.245 and p=0.124 respectively), FR variability (p=0.0907 and p=0.0793) and FR (p=0.105 and p=0.144). However, the strongest predictor of both change in MVC and CSA due to immobilisation was FR Variability.

**Figure 6:**
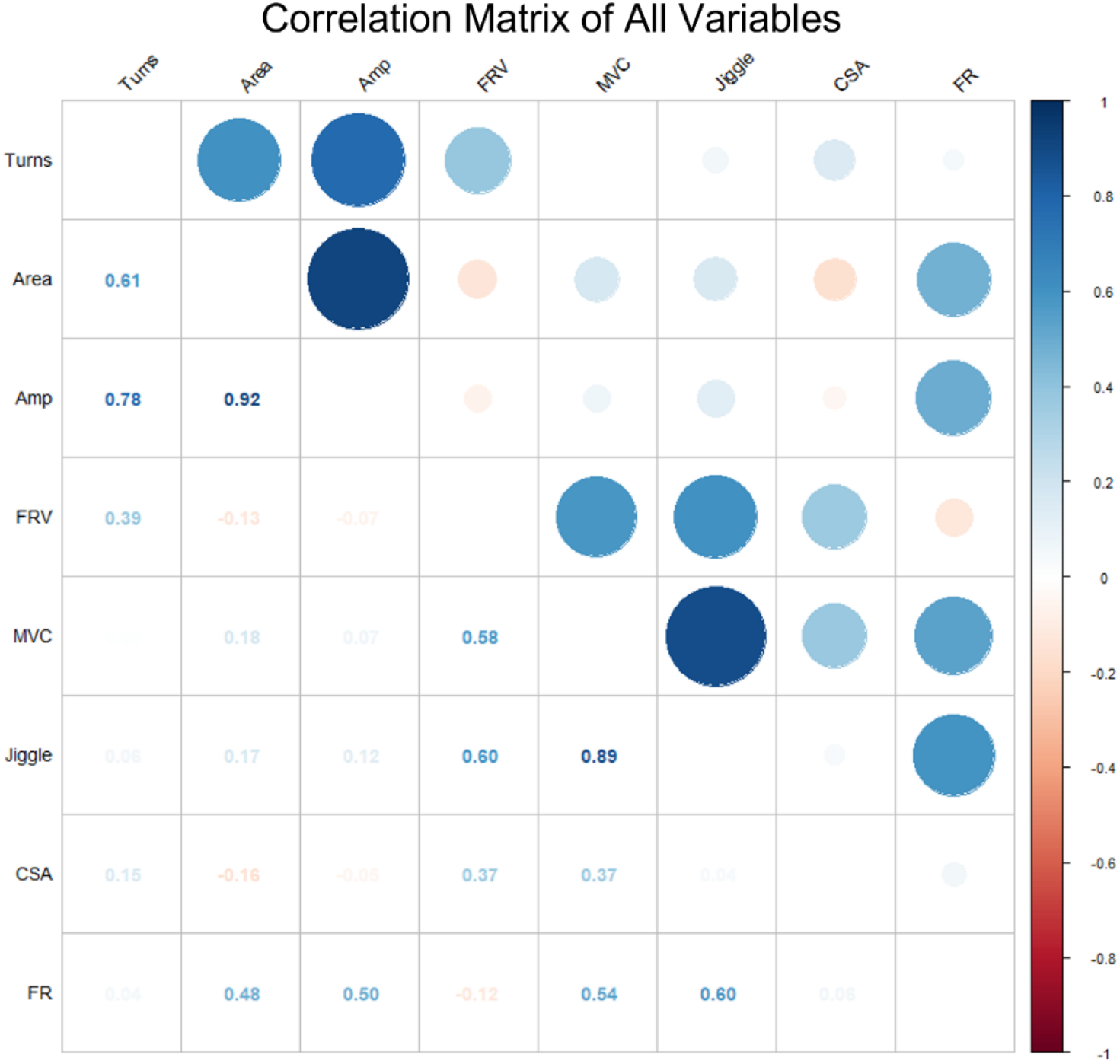
Correlation matrix of relationships between eight variables measured in this study. Amp; amplitude, FRV; firing rate variability, MVC; maximal voluntary contraction, CSA; cross-sectional area, FR; firing rate.

**Figure 7:**
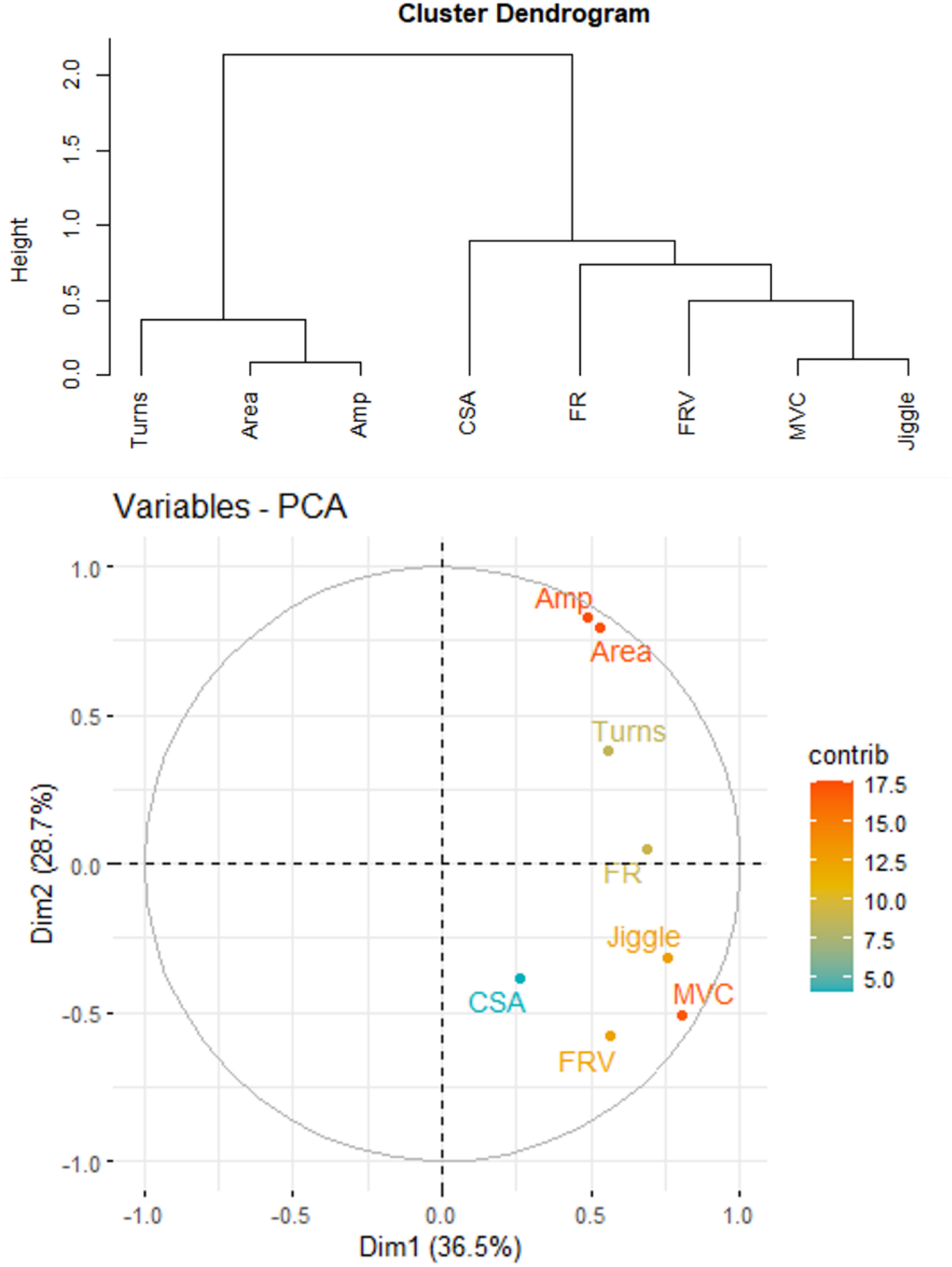
Cluster tree plot (upper) and accompanying principal component analysis plot (lower) to illustrate relationships between variables within the first two dimensions of variance between variables. Amp; amplitude, FRV; firing rate variability, MVC; maximal voluntary contraction, CSA; cross-sectional area, FR; firing rate, Dim; dimension.

## Discussion

This study has characterised the adaptations of individual VL MUs following 15 days unilateral lower-limb immobilisation, in immobilised and non-immobilised limbs. Our findings reveal a number of adaptations that are unique to the immobilised limb only, with the non-immobilised limb largely unaffected. Muscle of the immobilised leg became smaller and weaker, and individual MUPs became smaller and more complex. Motor units of the immobilised leg had greater NMJ transmission instability, with reduced FR. Notably, the majority of these decrements were still apparent when force levels were normalised to that achieved prior to immobilisation, highlighting impaired neural input to muscle as a prominent contributor to the observed reduction in muscle function.

The observed decline in muscle strength (−31%) in the immobilised limb exceeded the decline in muscle CSA (−15%), and this discordant finding supports several other studies employing a ULLS model; 14-day unilateral knee immobilisation with an ∼5% decrease in quadriceps CSA and a ∼25% decline in isometric strength (Glover *et al*., 2008), and following 14 days of limb cast immobilisation, an ∼8.5% decline in quadriceps CSA was observed alongside a ∼23% decline in muscle strength (Wall *et al*., 2014). Unilateral power output was reduced in the immobilised limb only. Bed rest studies have collectively shown a reduction in muscle performance of ∼3% per day, although this reduction in bed rest was seen to stabilise between day 5 and day 14, suggesting a sharp initial reduction (Di Girolamo *et al*., 2021) which may have also occurred in the present study. Furthermore, the control limb remained unaffected in terms of functional loss and does not support the concept of a negative cross-over effect on measures of neuromuscular function. Peak twitch force was unaltered when assessed with an involuntary contraction elicited from a single pulse to the femoral nerve, and likely reflects the lack of sensitivity of this method. Similarly, force steadiness was not affected by immobilisation at low or mid-range contraction intensities and indicates a preservation of basic motor control following disuse.

MUP size (area and amplitude) is reflective of the depolarisation of all fibres within a single MU, within the detection area of an indwelling electrode and is proportional to the size and number of fibres contributing to it. Area and amplitude were significantly decreased post-intervention at both 10 and 25% MVC contraction intensities in the immobilised leg only, and were unsurprisingly highly correlated. While greater MUP size increase is commonly observed in aged muscle reflecting MU remodelling and reinnervation of denervated muscle fibres (Jones *et al*., 2022), smaller MUPs differentiated sarcopenic from non-sarcopenic older men in this muscle group (Piasecki *et al*., 2018). The reduction observed here may indicate partial denervation of MU fibres as a result of disuse, or, it may reflect extensive fibre atrophy across MU fibres sampled at these contraction levels. MUP size reduction has also been observed in dystrophinopathies (Zalewska *et al*., 2013), where amplitude in the biceps brachii was significantly below reference values in patients with Duchene and Becker muscular dystrophies (Zalewska *et al*., 2004). All markers of MUP size increase with increasing contraction level, reflecting the recruitment of additional, larger MUs (Guo *et al*., 2022), and the decreases observed in the immobilised leg were apparent at 25% follow-up *and* baseline MVC. This finding highlights that reduction of force post-immobilisation cannot explain the decline in MUP size and is more likely related to a reduction in muscle fibre size and/or partial denervation/reinnervation of fibres.

Similar to MUP size, MUP complexity, defined as the number of MUP turns, was significantly greater in the immobilised leg only following the intervention, at both contraction intensities which is indicative of a greater temporal electrophysiological dispersion between individual fibres of the same MU (Stålberg & Sonoo, 1994; Piasecki *et al*., 2021*b*) as a result of an increased difference in conduction times along axonal branches and/or MU fibres. Increased MUP complexity has been observed in various myopathies which are suggestive of increased fibre diameter variability and also in neuropathies, expressing 75% greater turns than the control cohort, which is thought to be a product of the reinnervation and longer conduction times along axonal sprouts (Stewart *et al*., 1989). This has been reinforced specifically in myopathic conditions, as data from primarily the biceps brachii along with recordings of the vastus lateralis and gastrocnemius showed an 82% increase in polyphasic MUPs (Uncini *et al*., 1990). While much less severe, the changes seen following immobilisation may also follow a common pathway linked to denervation and reinnervation. This may suggest that some reinnervation has occurred following immobilisation, or alternatively that selective reduction in muscle fibre diameter has taken place, affecting MU fibres unequally resulting in more variable muscle fibre action potential propagation. Although needle insertions around the muscle motor point help to minimise the effects of variable muscle fibre conduction times and enable greater focus on axonal branch conduction variability, the specific motor endplate location remains an unknown *in vivo* and these effects cannot be completely excluded.

NMJ transmission instability, reported as NF MUP jiggle, was also significantly greater following the intervention in the immobilised leg only. NMJ structural stability was reported to decrease following 10-day bed rest, inferred following an increase in C-terminal agrin fragment (CAF) observed in serum at day 10 (Monti *et al*., 2021). It is suggested that CAF presence may be indicative of a dysregulation at the NMJ, reducing myoelectrical transmission reliability and therefore contractile activity in the muscle. However, it is not clear if this evidence represents active NMJ transmission as it is not directly measured during muscle contraction. Nevertheless, the *in vivo* methods applied here support the finding that disuse atrophy is accompanied by increased instability of transmission at the NMJ, possibly reflecting a partial denervation/reinnervation process. This process has been reported in healthy ageing (Hourigan *et al*., 2015; Piasecki *et al*., 2016*b*, 2021*a*) and diabetic neuropathy (Allen *et al*., 2015) corresponding with larger MUPs. Abnormally high NF MUP jiggle values have also been reported in the tibialis anterior of patients with chronic inflammatory demyelinating polyneuropathy (Gilmore *et al*., 2017) where an average increase of 90% compared to healthy controls was seen. This was associated with increased MUP area and complexity, also attributed to an incomplete reinnervation process. Herein, NMJ transmission instability was also greater when contraction intensity was normalised to baseline MVC following immobilisation, indicating that NMJ dysregulation is prevalent through a wider pool of motor units responsible for a broader range of contraction force.

The reduced MU FR reported here following immobilisation and loss of strength, may initially be explained by the lower absolute forces produced. However, this suppression of firing rate was also apparent at contractions normalised to baseline strength and clearly highlights it as a contributor to functional decrements. Ionotropic synaptic inputs and neuromodulation control the excitability of motoneurons, the latter of which is largely mediated by the amplitude of persistent inward currents (PICs) which act to amplify synaptic input and are proportional to the level of localised monoamine release (Heckman *et al*., 2008). Although difficult to quantify *in vivo*, it is possible monoamine levels decrease in response to reduced activity, yet it is unclear how this would influence the immobilised limb only. PIC amplitudes are also sensitive to inhibition (Hyngstrom *et al*., 2007; Mesquita *et al*., 2022), and recent RNAseq data highlight increased ligand-receptor interactions between muscle and dorsal root ganglion neurons following disuse, suggestive of an increased nociceptor sensitivity and susceptibility to pain (McFarland *et al*., 2022). As such, it is possible increased inhibition occurred in the immobilised limb only and suppressed PIC amplitudes and MU FR, similar to that believed to explain decreased FR following knee joint trauma (Nuccio *et al*., 2021). Reduction in firing rate variability at 10% MVC and 25% follow-up MVC may be viewed as a positive adaptation, however it more likely reflects the reduction in firing rate at lower absolute forces (Guo *et al*., 2022) as it was not apparent when force was normalised to baseline MVC.

Additional analysis into factors potentially related to the reduction in force and changes in neuromuscular parameters found a clustering of NMJ transmission instability, MU FR and FR variability with MVC and CSA. Following multivariate simple linear regression, no significant relationships were observed between either muscle size or strength with those neuromuscular parameters. This suggests that, in these data, no single variable fully explained the decline in muscle strength and size.

Since the average length of hospital stay in the United Kingdom as of 2018/19 was 4.5 days (Ewbank *et al*., 2020), future work in disuse should focus on the impact of such a short time frame on the neural input to muscle. Understanding these changes will provide a mechanistic basis on which to optimise rehabilitation protocols to counteract reduced muscle function.

### Limitations

A limitation to this study is that the contraction intensities at which motor units were sampled are of the low to mid-level and reveal nothing of adaptation to later recruited MUs. Knee extensor movements are not uniquely controlled by the VL, and although it may be a useful proxy for total quadriceps, we cannot rule out greater decrements in other muscles of this group. The current data are available in males only, and although we have highlighted similar differences across contractions in young males and females (Guo *et al*., 2022), there are sex-based differences in MU FR at normalised contraction levels which may respond differently to this intervention.

### Conclusion

These results support previous findings that unilateral short-term immobilisation of just 15 days leads to a decline in muscle strength unmatched by that in muscle size. Importantly, the current data highlight that this disconnect is explained by central and peripheral adaptations to neural input to muscle, evidenced by suppressed MU FR and increased instability of transmission at the NMJ, and identifies interventional targets for cases of muscle disuse.

